# Human Placenta-Derived Mesenchymal Stem Cells Stimulate Neuronal Regeneration By Promoting Axon Growth And Restoring Neuronal Activity

**DOI:** 10.1101/2023.10.26.561027

**Authors:** Elvira H. de Laorden, Diana Simón, Santiago Milla, María Portela-Lomba, Marian Mellén, Javier Sierra, Pedro de la Villa, María Teresa Moreno-Flores, Maite Iglesias

## Abstract

Cell therapy is a cutting-edge medical approach that involves the use of cells to treat various diseases and conditions. It harnesses the remarkable regenerative and reparative abilities of cells to restore or replace damaged tissues and promote healing.

In the last decades, mesenchymal stem cells (MSCs) have become the cornerstone of cellular therapy due to their unique characteristics. Specifically human placenta-derived mesenchymal stem cells (hPMSCs) are highlighted for their unique features, including ease to isolate, non-invasive techniques for large scale cell production, significant immunomodulatory capacity, and a high ability to migrate to injuries. Researchers are exploring innovative techniques to overcome the low regenerative capacity of Central Nervous System (CNS) neurons, with one promising avenue being the development of tailored mesenchymal stem cell therapies capable of promoting neural repair and recovery. In this context, we have evaluated hPMSCs as candidate for CNS lesion regeneration using a skillful co-culture model system. Indeed, we have demonstrated the hPMSCs ability to stimulate damaged rat-retina neurons regeneration by promoting axon growth and restoring neuronal activity both under normoxia and hypoxia conditions. With our model we have obtained neuronal regeneration values of 10-12% and axonal length per neuron rates of 19.99±0.77, μm/neuron. To assess whether the regenerative capabilities of hPMSCs are contact-dependent effects or it is mediated through paracrine mechanisms, we carried out transwell co-culture and conditioned medium experiments confirming the role of secreted factors in axonal regeneration. It was found that hPMSCs produce brain derived, nerve growth factors (BDNF, NGF) and Neurotrophin-3, involved in the process of neuronal regeneration and restoration of their physiological activity of neurons. The capability to access axonal physiology is crucial for studying information processing among neurons in healthy and diseased states. We confirm the success of our treatment using the patch clamp technique to study ionic currents in individual isolated living cells, confirming that in our model the regenerated neurons are electrophysiologically active.

## INTRODUCTION

Since the classic studies of Ramón y Cajal, the low regenerative capacity of the neurons of the Central Nervous System (CNS)(1) has been known. Since then, various strategies have been used to achieve neuronal regeneration, such as blocking axonal growth inhibitors or with cell therapy, using transplants of different cell types (2–18). For *in vivo* or future clinical studies, it would be essential to determine *in vitro* the neuroregenerative capacity of the cell populations that would be used in cell therapy10/26/23 2:10:00 PM. One of the best models to study *in vitro* and quantify cell-induced adult axonal regeneration is the co-culture of adult axotomized retinal ganglion cells (RGCs) with cells putatively capable of inducing axonal regeneration (olfactory ensheathing glia cells – OEGs, Schwann cells, astrocytes, etc.) (19–22). Our group has carried out studies in co-cultures of RGCs with OEG populations that have made possible to advance in the characterization of the molecular bases of the OEG-dependent regenerative capacity. We have shown that the ability of these cells to induce adult axonal regeneration in co-culture depends on several molecules (7,17,18): they secrete neurotrophic factors (23–25), they produce extracellular matrix proteases that contribute to degrading the perineuronal network that stabilizes the environment of adult neurons (26); produce proteases that stimulate axon regeneration (27), etc.

Stem cells are unspecialized cell precursors that are able to self-renew and differentiate into one or more specialized cell types in response to specific signals (28–31). In higher animals and based on their source, stem cells have been classified into two groups: on the one hand, embryonic stem cells (EScells), pluripotent, derived from the internal cell mass of the embryo at the blastocyst stage (7-14 days), that can generate all the different cell types in the body (32) and on the other hand, the organ-specific stem cells, derived from those, after many cell divisions, and which are multipotential; that is, they can originate the cells of a specific organ in the embryo, and also in the adult (33–36).

It has been shown, using animal models of spinal cord injury (SCI), that different stem cells, both embryonic and adult, undifferentiated or differentiated in culture with different protocols, are capable of promoting neuro-regeneration after injury, as well as functional recovery (37–39).

The regenerative potential of mesenchymal stem cells of different origins has been studied and contrasted by numerous research groups (40–42). The source from which organ-specific mesenchymal stem cells have traditionally been extracted is bone marrow. These cells can generate all cell types of the blood and the immune system (35), they can be grown both *in vitro* in the laboratory and *in vivo* (using animal models) and have been tested in tissue repair experiments (43,44). Transplantation of bone marrow-derived cells into SCI models has been reported to promote axonal regeneration, reduce lesion size, and improve functional *outcome* (45–53). The precise cell type within bone marrow responsible for these beneficial effects is not fully established but is thought to reside within the marrow stromal, corresponding to the MSC population (54,55). Additionally, transplantation after SCI of bone marrow MSCs gene-modified to secrete brain BDNF give way to increased corticospinal neurons survival in primary motor cortex as compared with the unmodified MSCs and promoted effects in locomotor recovery not observed with the control MSCs (56).

However, the use of bone marrow stem cells has some limitations due to the invasiveness of the extraction method and the decrease in their proliferation and differentiation capacity with the maintenance of cells in culture (30,57). Many researchers are working on finding an alternative to these cells that can be used in clinical applications. Placental tissue is a source of cells of great value in regenerative medicine because hPMSCs are easily isolated and can be expanded in culture using a suitable medium, they have great phenotypic plasticity, and in addition, due to the fact that the placenta is involved in the maintenance of fetal tolerance during pregnancy, they develop immunomodulatory properties of great importance in clinical applications based on cell therapy. These characteristics point to hPMSCs being suitable candidates for use in cell therapies for CNS injury (58–61).

For all these reasons, in this work we have, determined and quantified the ability of hPMSCs to induce adult axonal regeneration by using our model of co-culture with adult axotomized NGRs, under normoxic and hypoxic conditions. It is described that certain features of hPMSCs are stimulated when subjected to low oxygen concentrations in culture, including proliferation, migration, and neuroprotective potential (62–68). Furthermore, we understand that hypoxic conditions are representative of the physiological reality in some time course events after a CNS trauma or some neurological pathologies, such as a ischemic stroke. Therefore, we considered performing experiments simultaneously under normoxia and hypoxia conditions, which would allow us to compare the results in both environments.

Moreover, to assess whether the regenerative capabilities of hPMSCs are entirely contact-dependent effects or they are also mediated through paracrine mechanisms, we carried out Transwell co-culture and conditioned medium experiments confirming the role of secreted factors in RGCs’ axonal regeneration.

Finally, through the application of the patch clamp technique, we assessed the functional recovery of regenerated neurons, thus confirming their electrophysiological activity. This provides valuable insights into the realm of neural regeneration and its potential implications for treating neurological disorders.

## MATERIALS y METHODS

### Culture of hPMSCs

hPMSCs (Cellular Engineering Technologies, Coralville, USA, catalog number HMSC.AM-100) were grown in HGCM composed of high glucose Dulbecco’s modified Eagle’s medium (DMEM) (4.5 g/ l) (Gibco, Grand Island, New York, USA) supplemented with 10% fetal bovine serum (FBS) (Gibco, reference 10270106), an antibiotic and antifungal solution (Pen/Strep/Fung 10k/10k/25 µg, Lonza, Basel, Switzerland), 2 mM L-glutamine (Lonza), and basic fetal growth factor (hFGF basic) (Gibco). Cell cultures were incubated at 37 °C under normoxic conditions (5% CO_2_ and 20% O_2_) and hypoxic conditions (5% CO_2_ and 1% O_2_). The maintenance of hypoxia conditions in the cultures was monitored by assays by immunodetection of hypoxia inducible factor (HIF).

### Olfactory ensheathing cells (OECs)

OEGs line TS12 (69) was maintained in ME medium, composed of DMEM/F12 (Gibco) supplemented with 10% FBS, 2 mM glutamine (Lonza), 20 µg/mL pituitary extract (Gibco), 2 µM forskolin (Sigma) and an antibiotic and antifungal solution (Pen/Strep/Fung 10k/10k/25 µg, Lonza).

### Proliferation capacity of hPMSCs

To examine the proliferation capacity of the hPMSCs *in vitro*, the cumulative population doubling (PD) was calculated over 30 days (from passage 3 to 8). The hPMSCs were placed in triplicate into 10 cm^2^ multiwell dishes at a concentration of 10^4^ cells/cm^2^ and subcultured after 5 days at the same density. The cells were counted using a hemocytometer. The cumulative cell doubling of the cell populations was plotted against time in the culture to determine the growth kinetics of hPMSCs expansion. The number of population doubling was determined by counting the number of adherent cells at the start and end of each passage. The population doubling was calculated at every passage according to the equation: log_2_ (number of harvested cells/number of seeded cells). The finite population doubling was determined by the cumulative addition of the total numbers generated from each passage until the cells stopped dividing.

### Mesodermal differentiation of hPMSCs

For adipogenic differentiation hPMSCs were seeded at an inoculation density of 2.4×10^5^ cells/well in 6-well plates until they reached confluence and were then induced by three cycles of induction/maintenance with Differentiation media BulletKits – adipogenic (Lonza) according to the manufacturer’s instructions. Adipogenesis was assayed by staining of intracellular lipid droplets with Oil Red O (Sigma, St. Louis, Missouri, USA). The monolayer cultures were incubated with Oil Red O solution for 10 min at ambient temperature and examined by microscopy.

For osteogenic differentiation hPMSCs were seeded at an inoculation density of 3×10^4^ cells/well in 6-well plates and cultured in osteogenic induction medium Differentiation media BulletKit™ - osteogenic (Lonza) according to the manufacturer’s instructions. After 21 days in osteogenic induction medium, mineral deposits were observed by Alizarin Red (Sigma) staining. For Alizarin Red staining, the cells were fixed with 10% formalin for 5 min. The formalin was removed, and cells were washed twice with water. Then, the cells were incubated with Alizarin Red S staining solution (1.4%, pH 4.0; Sigma) for 20 min at ambient temperature and examined by microscopy.

For chondrogenic differentiation micromass cultures were generated by seeding 5 µL droplets of 1.6×10^7^ cells/mL cell solution in the center of multi-well plate wells. After cultivating micromass cultures for 2 hours StemPro® Chrondrogenesis differentiation medium (Gibco) was added. After 14 days of incubation, the micromasses were rinsed with PBS and fixed with 4% paraformaldehyde solution for 30 minutes. Chondrogenic differentiation was observed by Alcian Blue staining (Merck KGaA, Darmstadt, Germany) and examined by microscopy.

### Quantitative polymerase chain reaction (qPCR) for pluripotent stem cell markers

Total RNA from hPMSCs was isolated using RNeasy Mini Kit (QIAGEN, Hilden, Germany). Reverse transcription (RT) reactions were carried out on 250 ng of total RNA using the High-Capacity cDNA Reverse Transcription Kit” (Applied Biosystems, Waltham, Massachusetts, USA). RT-PCR analysis for pluripotency markers genes such as *OCT4*, *SOX* y *NANOG* was carried out. SDS Software (Applied Biosystems) was used to analyze the data.

### Flow cytometry analysis

To confirm MSCs phenotype of cells grown in normoxic and hypoxic conditions, cells were incubated with antibodies against the following human antigens: CD105, CD44, CD73, CD14, CD34, CD19, and CD45 and HLA-DR. After washing, cells were subjected to flow cytometry analysis using a flow cytometer (Miltenyi Biotech). The data were analyzed with MACSQuantify™ Software.

### Western blot assays

Protein lysates at different passages (from 3 to 5) and different oxygen concentration cultures of hPMSCs were obtained using lysis buffer (NaCl 150 mM, IGEPAL® 1%, DOC 0.5%, SDS 0.1%, Tris pH 8 50 mM) containing protease inhibitors (50X, Roche, Mannheim, Germany). Protein concentrations were determined using bicinchoninic acid assay (Thermo Fisher Scientific, Waltham, MA, USA). Equal amounts of protein (30 µg) were separated by 10% sodium dodecyl sulfate-polyacrylamide gel electrophoresis and transferred to polyvinylidene difluoride membranes (Roche). Membranes were blocked and incubated with primary antibodies at 4°C overnight: anti-BDNF (1:1,000, rabbit, Abcam, cat# ab108319), anti-NGF (1:1,000, rabbit, Abcam, cat# ab52918), anti-NT-3 (1:1,000, rabbit, Abcam, cat# ab75960), anti-HIF (1:1000, rabbit, Abcam, cat# ab51608) or anti-β-actin (1:1,000, mouse, Merck, cat# A5441). Next day, membranes were incubated with horseradish peroxidase-conjugated goat anti-rabbit (1:10,000, Cell Signaling Technology, cat# 7470) or sheep anti-mouse (1:10,000, Merck, cat# AC111P) secondary antibodies for 1 hour at ambient temperature. After applying ECL substrate (Amersham, Buckinghamshire, UK) the protein bands were chemiluminescently detected in ChemiDoc MP Imaging System (Bio-Rad) and the relative protein expression to β-actin was analyzed by Image J software.

### Axonal regeneration assays by direct co-culture

Axonal regeneration assays were performed and analyzed as previously described. Retinal tissue was extracted from 2-month-old rats, and then digested with papain using Worthington’s Papain System (20 units/ml; Worthington-Biochemical Corporation, Lakewood, NJ). Axotomized adult retinal neurons were plated onto hPMSCs (5×10^4^, 8×10^4^ and 10^5^) and OEG (1.1×10^5^) monolayers and the cultures were maintained in serum-free neurobasal medium (Gibco), supplemented with B-27 (Gibco) (NB-B27) for 96 hours in normoxia and hypoxia. Axotomized adult retinal neurons were also plated onto 10 mg/ml poly-L-lysine (Sigma) coated dishes as negative control. Co-cultures were fixed with 4% paraformaldehyde (PFA).

### Axonal regeneration assays by indirect co-culture

Axotomized retinal neurons were plated onto 10 mg/ml PLL (Sigma) coated dishes and cultured in NB-B27 medium. hPMSCs were seeded directly onto 24-well 0.4 µm Transwell inserts at a density of 10^4^ cells/cm^2^, cultured in NB-B27 medium and maintained in normoxia and hypoxia for 96 hours before fixing them with 4% PFA.

### Axonal regeneration assays using conditioned media

hPMSCs were seeded at a density of 10^4^ cells/cm^2^ in HGCM. When cultures reached approximately 90% confluence, hPMSCs were rinsed with PBS (PBS EDTA pH 7.5; Lonza) and maintained in NB-B27 medium for 96 hours. Conditioned medium was collected and centrifuged at 1,200 g for 5 min.

Axotomized neurons were plated as described above. Conditioned medium or equivalent volume of NB-B27 medium (control) was added to the neurons. The cells were incubated for 96 hours and fixed with 4% PFA.

Axonal regeneration events were thoroughly characterized and quantified as described previously (21,22,26). Co-cultures were analyzed by immunocytochemistry employing an antibody against a phosphorylated form of MAP1B and Neurofilament-H (NF-H) axonal proteins (SMI31; Sternberger Monoclonal Inc., Baltimore, MD; 1:500) and an antibody, 514, which recognizes high molecular weight microtubule-associated protein 2 (MAP2A,B; 1:400) (70). The percentage of retinal neurons with an axon was determined by counting the number of MAP2A,B positive neurons that bear a polarized neurite that can be labelled with the antibody against the phosphorylated forms of MAP1B and NF-H proteins. The samples were observed in an inverted fluorescent microscope with a 40X immersion objective (DMi8, Leica, Wetzlar, Germany) and a minimum of 20 fields (containing a minimum of 200 neurons) were taken randomly.

Two parameters were determined: percentage of neurons that have regenerated axons (number of neurons with an axon *versus* total number of counted neurons), and the mean axonal length of the regenerated axons (µm)/neuron (ratio between the total length of the regenerated axons and the total number of counted neurons). Fluorescence signals were analyzed with ImageJ software (Image J, NIH) and the length of the axons was determined using the Neuron J plugin.

#### Immunocytochemistry

Direct and indirect co-cultures, and cultures with conditioned media, were fixed with 4% PFA to perform immunocytochemistry analysis. Samples were permeabilized and blocked in blocking solution (1X PBS, 0.1% triton, 1% FBS) for 30 minutes at room temperature. Next, they were incubated with the corresponding primary antibody in blocking solution at the optimal dilution for each of them. The primary antibodies used were SMI31 and 514. Incubation was carried out at 4 °C overnight. After washing, the cells were incubated for another hour in a solution of PBS-TS containing the appropriate fluorescent secondary antibody, at its optimal dilution. Fluorescence-conjugated secondary antibodies used were Alexa Fluor 488 and Alexa Fluor 594, respectively.

Finally, the samples were washed with PBS and mounted with Flouromount (Southern Biotech, Birmingham, Alabama, USA). The labelled preparations were visualized using an inverted with fluorescent microscope with a 40X immersion objective (DMi8, Leica).

### Study of properties of regenerated axons

To detect the expression of mature synaptic vesicles and voltage-gated sodium channels (VGSCs), direct and indirect co-cultures were fixed, permeabilized and blocked as previously described. MAP2A,B (514) (27), SV2A (kind gift from Dr. Morcillo, University of Extremadura, Spain) and Nav1.1 α subunit (SCN1A) (Sigma, cat# S8809) primary antibodies were incubated at 4 °C overnight. Fluorescence-conjugated secondary antibodies used were Alexa Fluor 488 and Alexa Fluor 594.

### Brain-Derived Neurotrophic Factor (BDNF) and Nerve Growth Factor (NGF) quantification

BDNF and NGF levels in direct and indirect co-cultures, and in conditioned media, were evaluated using Human BDNF ELISA kit (Abcam, cat# ab1212166) and Human beta Nerve Growth Factor ELISA Kit (Abcam, cat# ab193760) following the manufacturer’s instructions.

### Electrophysiology

Electrophysiological recording of cell voltage and ionic currents were performed under the whole cell configuration of the platch clamp technique (Hamill, Marty, Neher, Sakmann, & Sigworth, 1981) using an Axopatch 200A amplifier (Axoninstruments, FosterCity, CA, USA). Borosilicate pipettes (1.2 mm outer diameter) with a tungsten internal filament, were made using a vertical pipette puller (Narishige mod. P88, Narishige, Tokyo, Japan). The inner diameter of the pipette after pulling was approximately 0.5 to 1 μm. Filling of the pipette was carried out with an intracellular solution containing (in mM): 10 NaCl, 110 KCl, 5 EGTA, 0.5 CaCl_2_, 1 MgCl_2_, and 10 glucose (pH 7.4). The electrical resistance of the pipettes was measured, in the range of 8 to 12 MΩ. The seal resistance was approximately 1 to 3 GΩ. Liquid junction potential was routinely corrected. The Ag-AgCl indifferent electrode was connected via an agarose bridge to the perfusion. The cell voltage was maintained at −80 mV and depolarizing pulses of 30 ms duration were applied in steps of 5 mV. Both the holding voltage and the pulses were generated using a personal computer connected to the CED plus (Cambridge Electronic Design Ltd, Cambridge, England). Data were sampled at 0.2–10 kHz after low-pass filtering with an appropriate cut-off for each sampling frequency. Data analysis was performed offline using a personal computer.

### Statistical analysis

Data are presented as the mean ± standard deviation (SD). Multiple comparisons, one-way ANOVA followed by Dunnet’s *post hoc* test was used to evaluate the differences in axonal regeneration. An alpha value of p ≤ 0.05 was used for statistical significance.

## RESULTS

### Culture and characterization of hPMSCs

Prior to evaluate neural regeneration potential of hPMSCs, we confirmed that the cells were in agreement with the criteria of International Society for Cellular Therapy published in 2006 (71).

Mesenchymal cells attached to cell culture plastic flasks and showed a typical spindle-type shape morphology when observed by inverted phase contrast microscopy (Figure 1A). No differences in morphology between oxygen conditions were detected.

**Figure 1.**
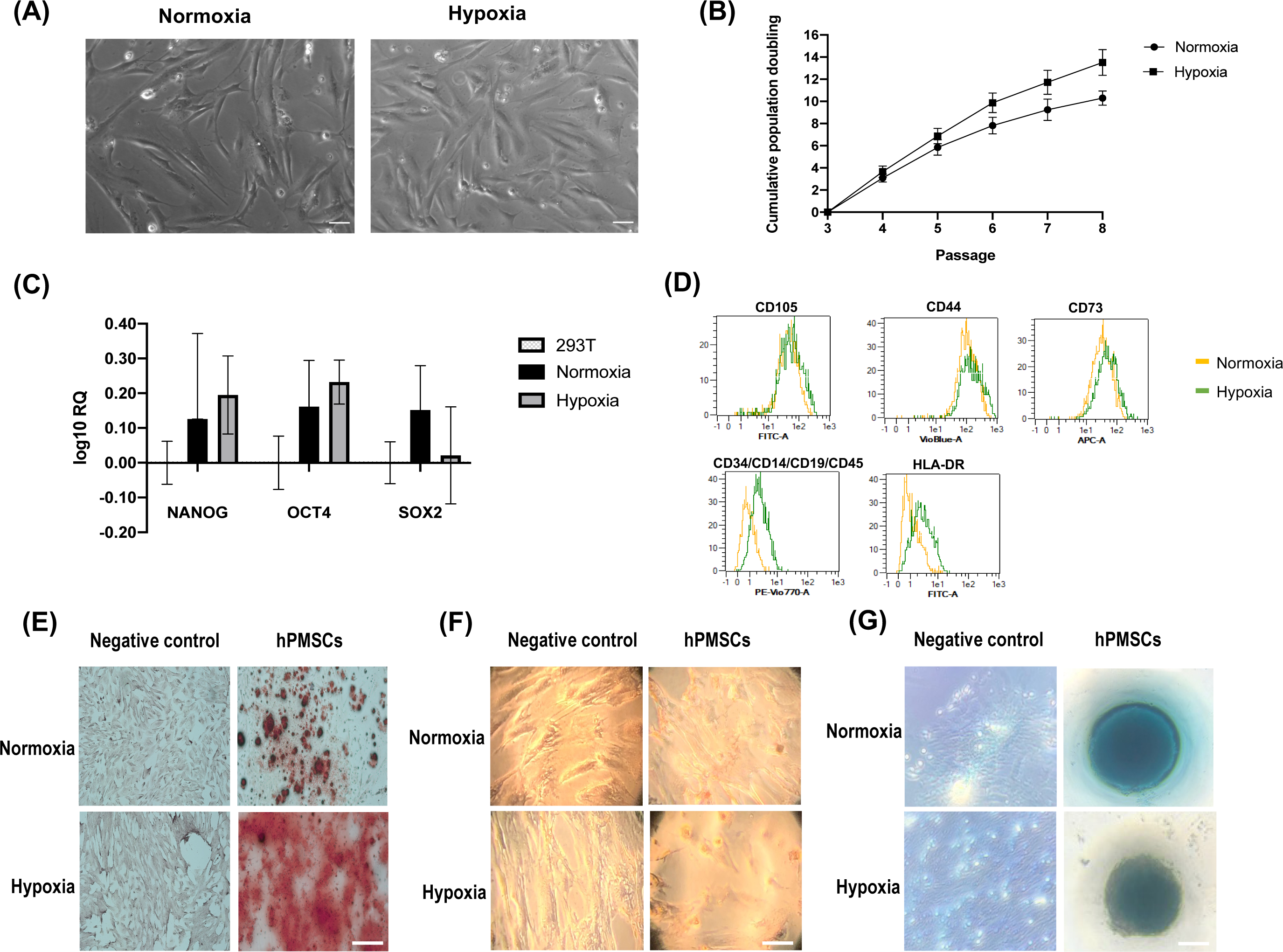
Characterization of hPMSCs. (A) Morphology at passage 3. Both normoxic and hypoxic cultures show highly homogeneous spindle-like shape. Scale bar: 500 µm (B) Proliferation capacity. Cumulative population doublings of hPMSCs grown under hypoxia (square) are higher than cultures grown in normoxia (dot). (C) Detection of pluripotency markers by RT-PCR shows that expression of *NANOG*, *OCT4* y *SOX2* is higher in hPMSCs than in differentiated cells. (D) Flow cytometry shows specific hMSCs marker expression pattern: positive for CD105, CD44, CD73 and negative for CD14, CD19, CD34, CD45 and HLA-DR. (E) Osteogenic differentiation was confirmed by Alizarin red staining of calcium deposits. (F) Adipogenesis differentiation was followed by Oil Red O staining of lipids vacuoles. (G) Alcian Blue staining of proteoglycans demonstrated chondrogenesis differentiation. Scale bar: 100μm.

The comparative growth kinetics of the hPMSCs grown in normoxia and hypoxia from passages 3 to 8 revealed that hypoxic cultures had a higher growth potential, measured as cumulative PDs (Figure 1B).

We then analyzed pluripotency markers expression (*NANOG*, *Oct-4* and *SOX2*) and observed an increased relative expression when compared to the fibroblast cell lineage 293T (Figure 1C).

To assess the expression of mesenchymal and hematopoietic markers an immunophenotypic analysis was performed. Flow cytometry results indicated that hPMSCs expressed typical immunophenotypic characteristics consistent with those of a mesenchymal lineage. Cells cultured in both oxygen conditions were positive for surface markers CD105, CD44 and CD73 and negative for CD34, CD14, CD 45, CD 19, and HLA-DR (Figure 1D).

We investigated the differentiation potential of hPMSCs by inducing them into osteoblasts, adipocytes and chondrocytes *in vitro*. Alizarin red staining revealed significant calcium deposition in treated cells, thus confirming osteogenesis (Figure 1E), Oil Red O-stained lipid drops were observed in differentiated hPMSCs indicating adipogenesis (Figure 1F), and the presence of proteoglycan staining with Alcian blue proved chondrogenesis (Figure 1G). Interestingly, hPMSCs grown under hypoxic conditions showed a substantial increase in calcium deposition when compared to hPMSCs grown in normoxia. Similarly, hypoxic hPMSCs cultures showed a higher number of lipids droplets. These results suggest that hypoxia promotes hPMSCs differentiation potential.

Taken together, these results confirm that the cells of this study were mesenchymal stem cells. In addition, these data reveal that low oxygen concentration has no significant *in vitro* effect on the morphology and phenotype profiles of hPMSCs but it can promote both hPMSCs proliferation and differentiation abilities.

### Neuroregenerative effect of hPMSCs

To evaluate hPMSCs neuroregenerative potential an *in vitro* regeneration assay was performed. All analyses were conducted on cells at passages 2-5, simultaneously cultured under normoxic and hypoxic conditions.

Different densities of hPMSCs (5×10^4^, 8×10^4^ and 10^5^) were co-cultured with axotomized 2-month-old rat retinal ganglion cells (RGCs) in both normoxia and hypoxia, for 96 hours. In parallel, an immortalized human olfactory ensheathing cell line (TS12) was used as a low regenerative capacity control (69). Wells treated with the substrate PLL with no cells attached were used as a negative control for RGCs axonal regeneration capacity. Neuroregeneration was detected by immunofluorescence using SMI31 antibody (against MAP1B and-NF-H) as an axonal marker and 514 antibody (against MAP2A,B) to observe the somatodendritic compartment (Figure 2A). Two parameters were then quantified: the percentage of neurons with an axon, and the mean axonal length per neuron (µm/neuron).

**Figure 2.**
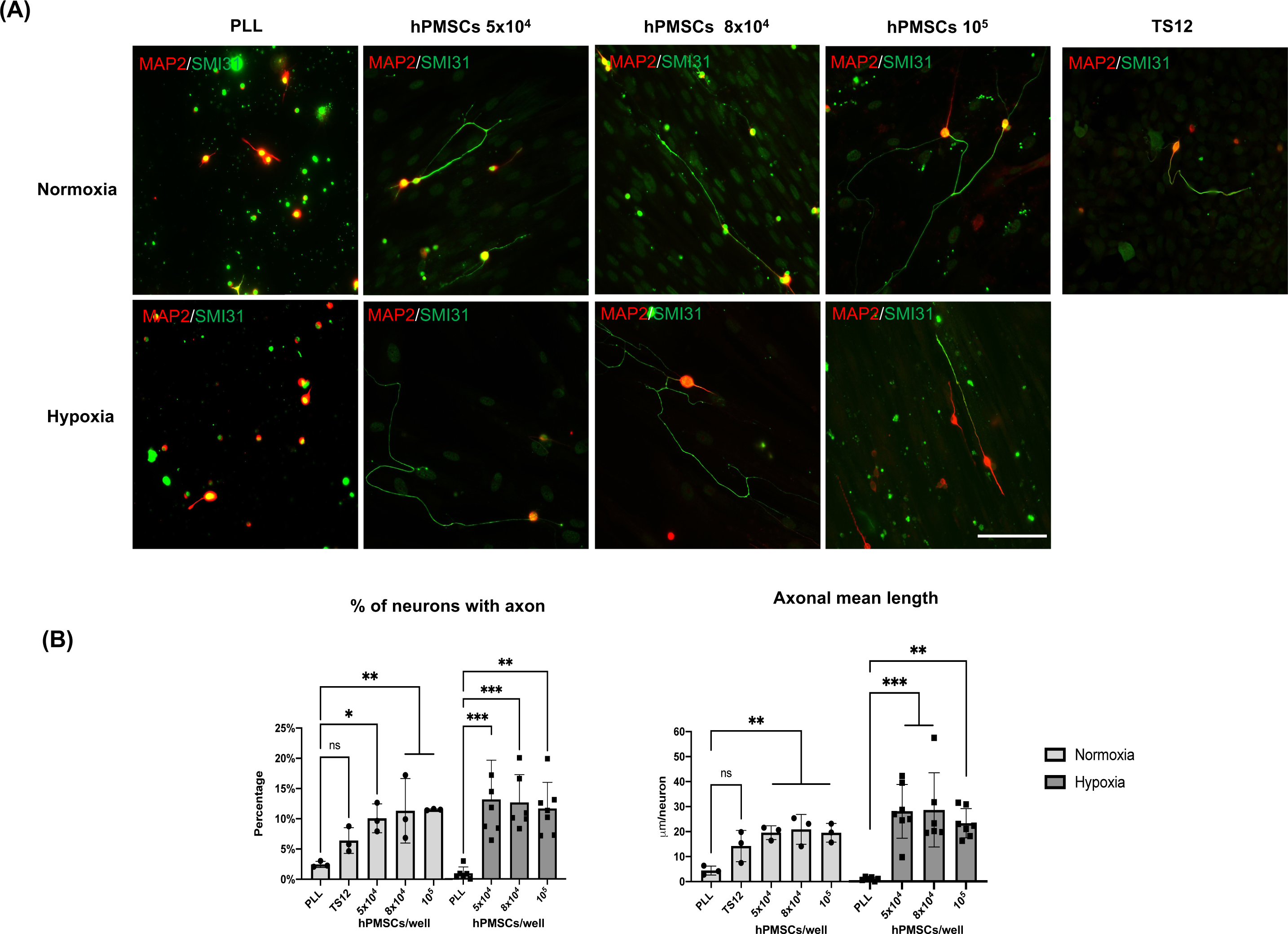
Neuroregenerative effect of hPMSCs. Axotomized adult RGCs were co-cultured with hPMSCs at different densities. TS12 cells were used as low regenerative control. Axotomized RGCs were also plated onto poly-L-lysine substrate (PLL) to measure the intrinsic neuroregenerative capacity. (A) Representative images of immunofluorescence of axon-specific Neurofilament-H protein (SMI31, green) and microtubule-associated protein 2 (MAP2A,B, red). Scale bar: 100 μm. (B) Axonal regeneration quantification: percentage of retinal neurons with an axon and axon mean length (μm/neuron). ANOVA and post-hoc Dunnet’s test. *p<0.05, **p<0.005 and ***p<0,001. (n=3 normoxia; n=6 hypoxia).

Axotomized RGCs showed a low regeneration capacity in normoxic PLL single cultures (2.5%±0.005 GCs with an axon). This capacity was lower in hypoxic atmosphere (1.04%±0.011 GCs with an axon) (Figure 2B).

However, the presence of hPMSCs significantly increased RGCs axonal regenerative capacity whereas TS12 cells did not (Figure 2B). The increment of RGCs with regenerated axon in the presence of hPMSCs was similar in both oxygen conditions, being slightly more efficient in hypoxia (14.58%±7.8, 14.01%±5.0, 12.90%±4.7 RGCs with an axon when co-cultured with 5×10^4^, 8×10^4^ and 10^5^ hPMSCs respectively) than in normoxia (10.08%±2.3, 11.33%±5.3, 11.52%±0.1 RGCs with an axon). Accordingly, mean axonal length in normoxia was 19.57±2.74, 20.88±5.96, 19.52±3.75 mm/neuron when RGCs were seeded with 5×10^4^, 8×10^4^ and 10^5^ hPMSCs respectively and 26.96±11.18, 25.80±14.28, 23.51±5.63 mm/neuron in hypoxia (Figure 2B). These subtle differences between hypoxia and normoxia were not statistically significant probably because the number of neurons per field analyzed is lower in hypoxia (7.7±0.59) than in normoxia (9.9±0.81) neurons per field. However, the axonal regeneration observed in hypoxia is significantly higher for both quantified parameters than its negative control (PLL), compared to that achieved under normoxia conditions.

Notably, these data demonstrate a high increment in axotomized RGCs regenerative capacity when cultured in the presence of hPMSCs.

### Neurotrophic factors expression by hPMSCs

Western blot experiments confirmed pro-BDNF, pro-NT3 and pro-NGF synthesis by hPMSCs when grown both in HGCM or NB-B27 media (Figure 3A,B) at any time point studied. After quantification by Image J (data not shown) we concluded that pro-NGF synthesis by hPMSCs was 2 times higher when grown in hypoxia relative to normoxia, in both types of culture media. Cell growth in hypoxia triggers the expression of HIF as it is described (72).

**Figure 3.**
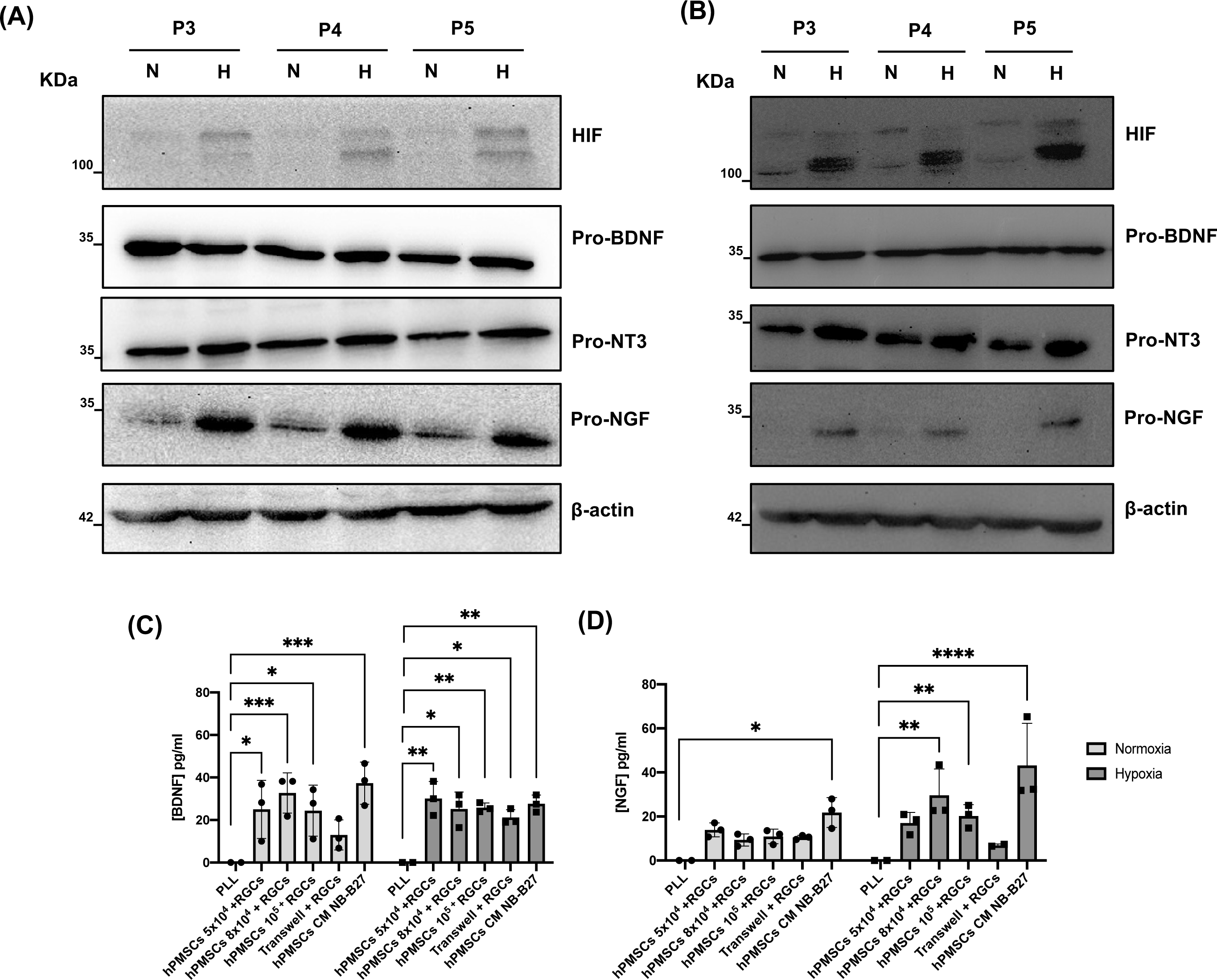
Neurotrophic factors expressed by hPMSCs. Western blotting was performed to analyze the synthesis of pro-BDNF (32 KDa), pro-NT3 (35 KDa), pro-NGF (32 KDa) in hPMSCs cultures at passage 3 (P3), passage 4 (P4), passage 5 (P5), grown in normoxia (N) and hypoxia (H). hPMSCs were cultured in (A) DMEM high glucose (4,5 gr/L) or (B) NB-B27 media. Maintenance of hypoxic conditions was monitored by immunodetection of hypoxia-inducible factor (HIF). Relative protein levels were controlled with β-actin expression. Secreted BDNF (C) and NGF (D) were measured using ELISA-based assays. ANOVA and post-hoc Dunnet’s test. *p<0.05, **p<0.005, ***p<0,001 and ****p<0,0001.

We confirmed the secretion of neurotrophic factors by hPMSCs. The concentration of secreted BDNF by hPMSCs (5×10^4^, 8×10^4^ and 10^5^ cells/plate) when maintained in co-culture with RGCs in normoxia is 25±13.68, 32.74±9.49 and 24.4±11.99 pg/ml, respectively. In hypoxia, secreted BDNF achieves a concentration of 30.12±8.04, 25.24±7.92 and 25.83±2.18 pg/ml. The concentrations are lower in indirect Transwell co-cultures (12.97±7.09 pg/ml in normoxia, 21.19±3.61 pg/ml in hypoxia). hPMSCs grown in NB-B27 medium (hPMSCs CM NB-B27) in absence of RGCs secreted a higher concentration of BDNF as a mean, but the difference was not significant attending culture conditions (Figura 3C).

Secreted NGF concentration is slightly higher in hypoxia (17.06±4.76, 29.66±11.93 and 20.23±5.15 pg/ml in 5×10^4^, 8×10^4^ and 10^5^ hPMSCs+RGCs co-cultures, respectively) than in normoxia (13.93±3.17, 9.33±2.75 and 10.92±3.30 pg/ml) (Figure 3D).

### Neuroregenerative effect of hPMSCs by indirect co-culture and conditioned medium collected from hPMSCs

Our previous results show that hPMSCs provided a permissive substrate that allowed axon regeneration and elongation in axotomized RGCs linked to neurotrophic pro-factors synthesized by hPMSCs. In order to evaluate the paracrine effect of neurotrophic factors secreted by hPMSCs two kind of *in vitro* indirect co-culture assays were performed.

For indirect co-culture assay: axotomized RGCs were plated onto PLL coated dishes. hPMSCs cell suspension (10^4^ cells/cm^2^) in NB-B27 medium was seeded onto the 0.4 µm Transwell inserts. These indirect co-cultures were performed in both normoxia and hypoxia. For secretome activity assay: NB-B27 medium was conditioned by hPMSCs in normoxia or hypoxia and later added to axotomized RGCs cultures on PLL (Figure 4A). Axonal regeneration was characterized and quantified as described previously (Figure 4B).

**Figure 4.**
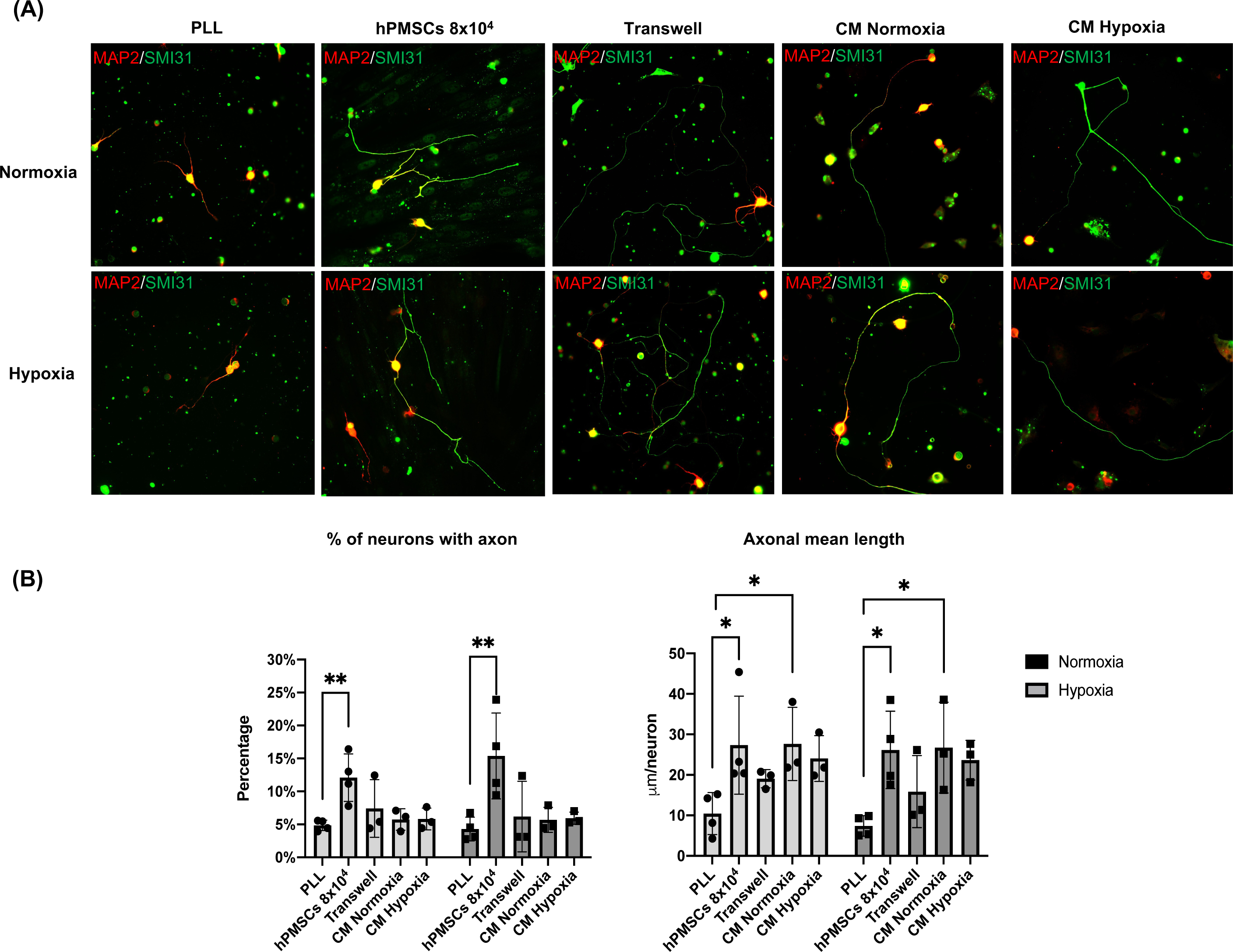
Paracrine effect of neurotrophic factors secreted by hPMSCs. Axotomized adult retinal neurons were plated onto poly-L-lysine substrate (PLL) to measure the basal RGCs neuroregenerative capacity. Co-cultures with hPMSCs (8×10^4^ cells) were used as positive control. Indirect co-cultures of axotomized neurons and hPMSCs were performed using Transwell inserts. Conditioned media both in normoxic (CM Normoxia) and hypoxic (CM Hypoxia) atmosphere were added to axotomized retinal neuron in culture. (A) After 96 hours of culture, cells were fixed and immunostained with antibodies SMI-31 (MAP1B/NF-H axonal marker, green) and 514 (MAP2A,B somatodendritic marker, red). Representative images are shown. (B) Axonal regeneration quantification: percentage of retinal neurons with an axon and mean axonal length per neuron (μm/neuron). ANOVA and post-hoc Dunnet’s test. *p<0.05 and **p<0.005. (n=3 normoxia; n=3 hypoxia).

Results demonstrated the paracrine effect of hPMSCs on RGCs neuroregeneration. In indirect Transwell co-cultures, the percentage of RGCs with a regenerated axon is very similar to control cultures (PLL) in normoxia (7.41%±4.3) and hypoxia (6.19%±5.3). However, mean axonal length per neuron achieved is more than 2 times higher in both conditions (19.07±2.24 mm/neuron in normoxia; 15.87±8.9 mm /neuron in hypoxia).

Similar results were obtained when axotomized RGCs were cultured with hPMSCs CM. Under those conditions, although the percentage of RGC with a regenerated axon is similar to the control (PLL), the mean axonal length per neuron is higher in both normoxia and hypoxia. No differences were observed in the effect on regeneration when CM were conditioned in either normoxic or hypoxic conditions (Figure 4B).

These results strongly suggest that hPMSCs function as a biological substrate that increase the capacity of RGCs to regenerate their axons, partly through a paracrine mechanism.

### Study of functional properties of regenerated axons

After neuroregeneration potential of hPMSCs was confirmed, we further analyzed the developmental stage of the regenerated axons by the detection of mature synaptic vesicles and voltage-gated sodium channels (VGSCs). Additionally, we explore the axonal complete functionality by measuring their electrophysiological capacity.

SV2A is a synaptic vesicle membrane glycoprotein expressed exclusively in neurons and endocrine cells essential in neurotransmitter release (73). To test the production of mature synaptic vesicles, SV2A protein expression was detected in regenerated axons of RGCs after hPMSCs co-culture. We observed several SV2A positive *puncti* throughout RGCs regenerated axons, with similar size and shape as previously described (74) (Figure 5A). Those images demonstrated the presence of mature synaptic vesicles along those axons, suggesting their ability to release neurotransmitters in both oxygen conditions.

**Figure 5.**
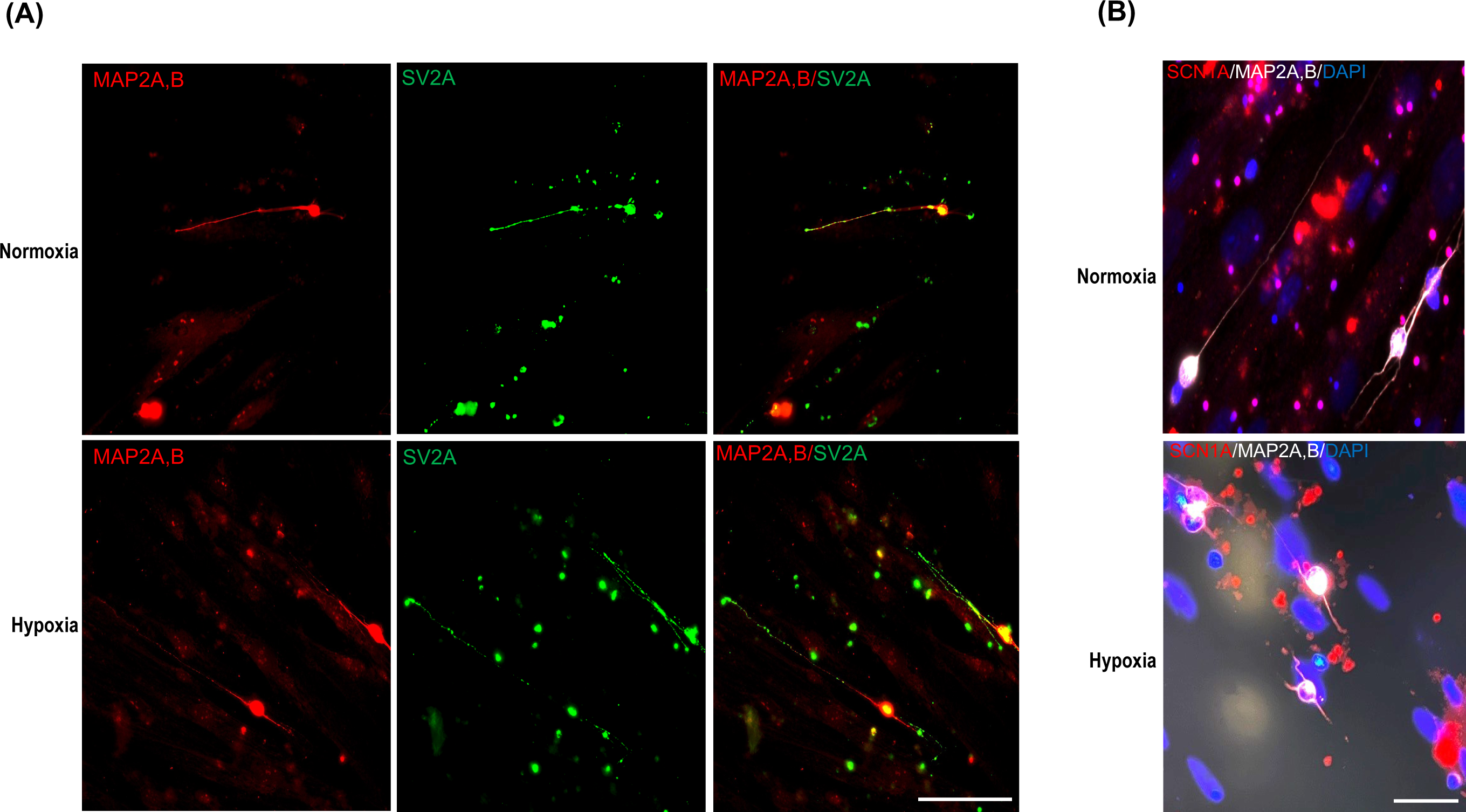
Study of properties of regenerated axons. Axotomized adult retinal neurons were co-cultured with 8×10^4^ hPMSCs/well. (A) After 96 hours of culture cells were fixed and immunostained with antibodies 514, against MAP2A,B (somatodendritic marker, red) and anti-SV2A, against synaptic vesicle membrane glycoprotein 2 (green). Representative images are shown. (B) Representative immunofluorescence of voltage-gated sodium channel Nav1.1 α subunit (SCN1A, red), 514 against MAP2A,B (white) and nuclear staining with DAPI (blue). Scale bar panel A: 100μm; scale bar panel B: 50μm.

VGSCs are responsible of the initiation and propagation of potentials in excitable cells. VGSCs Nav1.1 α subunit (SCN1A) is expressed predominantly in cell bodies and dendrites and participate in generation of both somatodendritic and axonal action potentials (75,76) Immunostaining of SCN1A in hPMSCs-induced regenerated RGCs axons revealed in both conditions the presence of VGSCs in RGCs bodies, dendrites, and initial part of the axons (co-stained with MAP2A,B) suggesting their ability to generate action potentials. (Figure 5B)

Taken together, these results confirm that RGCs axons, regenerated by hPMSCs induction, may have developed the subcellular and molecular capacity to be fully functional.

Once the neuroregenerative potential of hPMSCs and the expression of VGSC were confirmed, we proceeded to functionally explore the electrical activity of the regenerated neurons. For this purpose, we recorded the ionic currents generated by the cells maintained in normoxic and hypoxic conditions, by using the patch clamp technique, in “whole cell” configuration.

In a series of voltage clamping experiments carried out in regenerated cells, the ionic currents activated by cell depolarization were recorded. Depolarizing pulses were applied from a membrane potential of -80 mV in 5 mV step. Figure 6 shows the depolarization-activated currents recorded in a normoxic condition sample (Figure 6A), as well as in a hypoxic condition sample (Figure 6B). In both conditions, it is possible to observe the transient sodium current (I_Na_), followed by the sustained potassium current (I_K_). These experiments, carried out in a group of regenerated axons in normoxia (n=8) and regenerated axons in hypoxia (n=7), allowed us to observe that the amplitude of the sodium currents recorded from regenerated cells in hypoxia was significantly lower than the current from the normoxic regenerated cells (Figure 6C). Similarly, the amplitude of potassium currents showed less amplitude in the hypoxic group than in the normoxic group (Figure 6D). Despite the observed differences, both cell types were able to generate action potentials when recorded in current clamp mode (Figure 6E,F). Therefore, our functional experiments demonstrate that the regenerated cells are active from an electrophysiological point of view.

**Figure 6.**
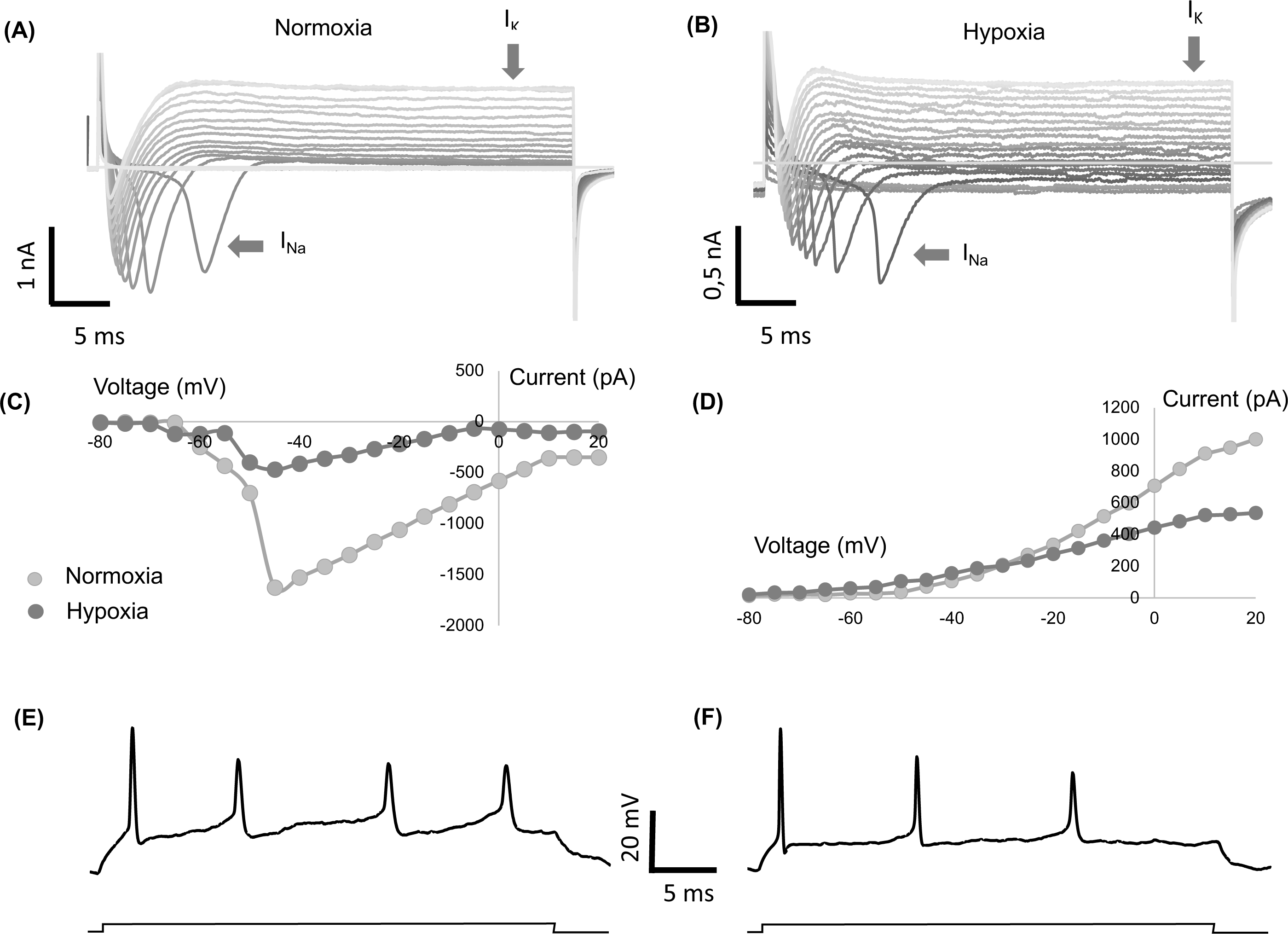
Study of electrical properties of regenerated axons. Axotomized adult retinal neurons were co-cultured with 8×10^4^ hPMSCs/well and recorded under voltage/current clamp conditions. (A,B) Representative whole cell current recordings from cultured cells in response to voltage pulses of 5 mV increasing step from a holding voltage of -80 mV in conditions of normoxia (A) and hypoxia (B). (C,D) Current voltage relationship of the sodium currents (C) and potassium currents (D) averaged from a total of 8 cells in normoxic conditions and 7 cells in hypoxic conditions. (E,F) Representative whole cell voltage recordings from cells shown in A and B in response to a current pulse of 0,1 nA.

## DISCUSSION

Cell therapy is a state-of-the-art medical paradigm that encompasses the utilization of cellular entities to address diverse pathological conditions. It capitalizes on the exceptional regenerative and reparative proficiencies intrinsic to cells, facilitating the re-establishment or substitution of impaired tissues and fostering the recuperative process. Cellular therapy represents a paradigm-shifting prospect poised to redefine the healthcare landscape covering a spectrum of disorders including those affecting the nervous system. This pioneering domain encapsulates diverse modalities of cell-mediated interventions, prominently featuring stem cell therapy (58,77–81).

Stem cell-based regenerative medicine is increasingly seen as a compelling approach for developing therapies. Among the most coveted cell sources, MSCs stand out due to their distinctive attributes. MSCs orchestrate tissue development, maintenance, and repair, and are useful for regenerative therapies to treat degenerative diseases and other clinical conditions (82–85).

Placenta-derived mesenchymal cells (hPMSCs) are a notable MSC source. They offer accessibility without invasiveness, possess strong immunomodulatory properties, high proliferative and differentiation potential, have the ability to migrate to injury sites *in vivo* and are ethically sound and versatile. For all these reasons, hPMSCs hold promise for regenerative medicine and better patient outcomes (59,86–89).

Recognizing the significant potential of these cells as therapy, our research group has directed its efforts towards further exploration and enhanced understanding of hPMSCs’ potential as cellular strategy to achieve neuronal regeneration.

As a preliminary and crucial step in the study we ensured that the properties of the cells in culture remained consistent across passages. Ensuring that cells maintain their intended characteristics over time and culture conditions is crucial for the reliability and reproducibility of experiments and future therapeutic interventions.

The cells, maintained in culture from passages 2-8, grew adherent to plate and exhibited a typical spindle-shaped morphology as observed under inverted microscopy. No observable alterations in cell morphology were detected by optical microscopy. Following the examination of pluripotency markers (NANOG, Oct-4, and SOX2), it became apparent that the cells preserved their pluripotent characteristics. An immunophenotypic evaluation using flow cytometry assay was performed to evaluate the expression of mesenchymal and hematopoietic markers. Results confirmed that the cells exhibited immunophenotypic characteristics consistent with those of a mesenchymal lineage when maintained in culture. Specifically, the cells were positive for the surface markers CD105, CD44, and CD73, while they were negative for CD14, CD34, CD19, CD45 and HLA-DR. The cellular differentiation capacity was also verified. In summary, our data confirm that the cells maintain their mesenchymal characteristics throughout the culture procedure and treatments (71).

In relation to both environments to which the hPMSCs were exposed, growth kinetics assays revealed that hypoxic cultures exhibited a markedly higher cumulative population doubling potential. In addition to that our results strongly suggest that hypoxia improves the differentiation potential of these cells.

After thoroughly and rigorously defining and characterizing our hPMSCs, we tackled the experiments to investigate the potential of these cells in the process of adult CNS axotomized neurons axonal regeneration and restoration of neuronal activity. To address this goal, we used an *in vitro* co-culture model, previously established by Dr. Moreno-Flores’s team. Our group considered applying a similar approach to track and characterize the potential of hPMSCs in an axonal regeneration process. Specifically, hPMSCs and axotomized 2-month-old rat retinal ganglion cells (RGCs) shared the culture plate for 96 hours and percentage of neurons with axon and mean axonal length per neuron were quantified (21).

With our experimental approach, we achieved success getting 5 to 13-fold increase in the percentage of neurons with regenerated axons respect to RGCs in PLL. Moreover, the mean axonal length increased 4 and 24 times under normoxic and hypoxic conditions respectively. This result settled a very significant and promising result.

Numerous researchers have focused their efforts on exploring potential therapeutic strategies rooted in stem cell therapy for addressing conditions linked to neuronal degeneration or dysfunction. Nonetheless, while the predominant approach in these studies revolves around assessing the differentiation capacity of MSCs to support damaged cells, they ultimately conclude that the observed effects are mediated through paracrine signaling. A prevailing consensus among research collectives underscores the critical role of stem cells in facilitating the regeneration of injured nerves, primarily through the secretion of neurotrophic factors, including BDNF, NGF, NT-3 (23,59,61,79,80,90–95).

In line with previous studies, we confirmed that hPMSCs express the neurotrophic factors BDNF, NT3 and NGF. The western-blot results referred to the pro-factors expression indicate that pro-BDNF and pro NT-3 expression level were homogeneous along time in culture and with the different oxygen concentrations. However, the expression of the pro-NGF increased markedly when the cells were maintained in hypoxia.

Since the pro-factors should be cleaved intracellularly or extracellularly to generate their mature forms, we wanted to determine BDNF and NGF present in the conditioned media. It has been widely proven the role of BDNF as a key mediator in axonal regeneration by stimulating axonal growth and facilitating synaptic reorganization. It is demonstrated the role of BDNF in axonal regeneration of RGCs in co-culture with primary and immortalized rat OEG (23). In our system, we have found that hPMSCs (5×10^4^ – 10^5^) release BDNF in concentration between 25±4.65 and 32±2.66 pg/ml with no difference when we compared secretion by cells maintained in normoxia and hypoxia conditions.

NGF plays a crucial role in promoting the growth and survival of sensory neurons. Both the expression and the secretion of this factor appear to be hypoxia dependent. We have observed a twofold increase when comparing cultures maintained under hypoxia respect to normoxia.

It was of great interest to us to verify whether the RCGs regenerative effect of hPMSCs was due to a cell-to-cell contact, proximity-derived or a paracrine effect. To determine whether the previously tested effect was necessarily linked to a cell-cell contact we conducted indirect co-culture experiments, and secretome activity assays. The percentage of neurons exhibiting axonal regeneration in presence of conditioned media would quantify the contribution to this regeneration attributable to a neurotrophic impact of the hPMSCs. Furthermore, the mean axonal length per neuron provides an indicator for assessing the extent of axonal growth resulting from that neurotrophic influence. In addition to identified neurotrophic factors, other uncharacterized molecules secreted into the media may be playing a role in these regenerative parameters.

Significantly, in this experimental framework, we observed a reduction in the number of neurons exhibiting regenerated axons compared to the co-culture experiments involving RGCs and hPMSCs contact. Indeed, the number of regenerated RGCs closely mirrored the results obtained in RGC cultures without hPMSCs. Nevertheless, the paracrine impact of hPMSCs on axon length per neuron, under both normoxic and hypoxic conditions, exhibited an approximately three to fourfold increase when compared to the control RGCs cultured on PLL plates. These results suggest that hPMSCs play a key role as biological substrate strengthening the capacity of RGCs to regenerate their axons, partly through a paracrine mechanism. Additionally, we have shown that capacity to promote the highest axonal regeneration in this experimental paradigm depends on contact between the adult RGCs and the hPSMCs, as demonstrated for other regenerative cells, such as OEGs (27,90,96).

We also wanted to confirm the regenerated RGCs functionality. Synaptic vesicles were localized distributed along the regenerated axons, suggesting their capacity to release neurotransmitters. In addition to that we assessed the regenerated axons’ ability to generate action potentials by characterizing the presence and activity of voltage-gated sodium channels (VGSCs). VGSCs play a pivotal role in the initiation and propagation of action potentials in neurons. Our findings demonstrate that the density of these channels is notably concentrated in the axon initial segment, a specialized region where action potentials are significantly greater than in the soma contributing to the distinctive electrical properties of this region. Based on our results, we can affirm that RGCs possess the molecular framework required for functionality.

Voltage-clamp experiments are a common method in neuroscience to measure the ionic currents in neurons. By controlling the membrane potential of the neuron, it is possible to measure the flow of ions in and out of the neuron during depolarization, providing valuable information about the neuron’s function and health (97).

In our experiments, we have confirmed the expression of VGSCs in the regenerated neurons. Moreover, we proceeded to compare the ionic currents generated by cells maintained in normoxia or hypoxia culture. Our findings demonstrate that regenerated neurons, under both environmental conditions, manifest transient sodium currents (I_Na_) and sustained potassium currents (I_K_) upon depolarization. Nevertheless, the amplitude of both sodium and potassium currents was notably reduced in the hypoxia-regenerated cells compared to those maintained under normoxic conditions, which could be due to the number and/or type of channels. In spite of these disparities, both cell types exhibited the capability to initiate action potentials, thereby affirming the electrophysiological activity of the regenerated cells representing without a doubt a biological breakthrough.

The study of the remarkably properties of hPMSCs, and the comprehensive understanding of all the molecules contributing to their ability to foster adult axonal regeneration undoubtedly warrant further investigation and approaches.

## Author Contributions

EHL, DS, MP-L and JS contributed to the performance of co-culture experiments. SM and PDV designed electrophysiology assays and analyzed the data. EHL and SM performed patch clamp assays. EHL extracted and analyzed the data. MI and MM obtained funding for the study. MI, MTM-F and EHL designed research. MI, EHL and MTM-F wrote the manuscript with inputs from all authors. All authors read and approved the final manuscript.

## Acknowledgments

We would like to acknowledge Prof. Morcillo (Universidad de Extremadura, Spain) for providing SV2A antibody. Dra. Pilar Martin-Duque for her support to our research.

## BIBLIOGRAPHY

1. Ramon y Cajal, S. Degeneration and Regeneration of the Nervous System, translated by May, R.M. London: Oxford University Press; 1928.

2. Horner,P.J. & Gage,F.H. Regenerating the damaged central nervous system. Nature. 2000;407:963–70.

3. Jones LL, Oudega M, Bunge MB, Tuszynski MH. Neurotrophic factors, cellular bridges and gene therapy for spinal cord injury. J Physiol. 2001;533(Pt 1):83–9.

4. Merkler D, Metz GA, Raineteau O, Dietz V, Schwab ME, Fouad K. Locomotor recovery in spinal cord-injured rats treated with an antibody neutralizing the myelin-associated neurite growth inhibitor Nogo-A. J Neurosci. 2001;21(10):3665–73.

5. Raisman G. Olfactory ensheathing cells - another miracle cure for spinal cord injury? Nat Rev Neurosci. 2001;2(5):369–75.

6. Bradbury EJ, Moon LDF, Popat RJ, King VR, Bennett GS, Patel PN, et al. Chondroitinase ABC promotes functional recovery after spinal cord injury. Nature. 2002;416(6881):636–40.

7. Moreno-Flores M, Díaz-Nido J, Wandosell F, Avila J. Olfactory Ensheathing Glia: Drivers of Axonal Regeneration in the Central Nervous System? J Biomed Biotechnol. 2002;2(1):37–43.

8. Okano H. Stem cell biology of the central nervous system. J Neurosci Res. 2002;69(6):698–707.

9. David S, Lacroix S. Molecular approaches to spinal cord repair. Annu Rev Neurosci. 2003;26:411–40.

10. Woolf CJ. No Nogo: Now Where to Go? Neuron. 2003;38(2):153–6.

11. Barnett S, Chang L. Barnett, S. C. & Chang, L. Olfactory ensheathing cells and CNS repair: going solo or in need of a friend? Trends Neurosci. 27, 54-60. Trends in neurosciences. 2004;27:54-60.

12. Nikulina E, Tidwell JL, Dai HN, Bregman BS, Filbin MT. The phosphodiesterase inhibitor rolipram delivered after a spinal cord lesion promotes axonal regeneration and functional recovery. Proc Natl Acad Sci U S A. 2004;101(23):8786–90.

13. Pearse DD, Pereira FC, Marcillo AE, Bates ML, Berrocal YA, Filbin MT, et al. cAMP and Schwann cells promote axonal growth and functional recovery after spinal cord injury. Nat Med. 2004;10(6):610–6.

14. Sandvig A, Berry M, Barrett LB, Butt A, Logan A. Myelin-, reactive glia-, and scar-derived CNS axon growth inhibitors: expression, receptor signaling, and correlation with axon regeneration. Glia. 2004;46(3):225–51.

15. Silver J, Miller JH. Regeneration beyond the glial scar. Nat Rev Neurosci. 2004;5(2):146–56.

16. Sivasankaran R, Pei J, Wang KC, Zhang YP, Shields CB, Xu XM, et al. PKC mediates inhibitory effects of myelin and chondroitin sulfate proteoglycans on axonal regeneration. Nat Neurosci. 2004;7(3):261–8.

17. Moreno-Flores MT, Avila J. The quest to repair the damaged spinal cord. Recent Pat CNS Drug Discov. 2006;1(1):55–63.

18. Gómez RM, Sánchez MY, Portela-Lomba M, Ghotme K, Barreto GE, Sierra J, et al. Cell therapy for spinal cord injury with olfactory ensheathing glia cells (OECs). Glia. 2018;66(7):1267–301.

19. García-Escudero V, García-Gómez A, Gargini R, Martín-Bermejo MJ, Langa E, de Yébenes JG, et al. Prevention of senescence progression in reversibly immortalized human ensheathing glia permits their survival after deimmortalization. Mol Ther. 2010;18(2):394–403.

20. Lim F, Martín-Bermejo MJ, García-Escudero V, Gallego-Hernández MT, García-Gómez A, Rábano A, et al. Reversibly immortalized human olfactory ensheathing glia from an elderly donor maintain neuroregenerative capacity. Glia. 2010;58(5):546–58.

21. Portela-Lomba M, Simón D, Russo C, Sierra J, Moreno-Flores MT. Coculture of Axotomized Rat Retinal Ganglion Neurons with Olfactory Ensheathing Glia, as an In Vitro Model of Adult Axonal Regeneration. J Vis Exp. 2020;(165).

22. Moreno-Flores MT, Lim F, Martín-Bermejo MJ, Díaz-Nido J, Avila J, Wandosell F. Immortalized olfactory ensheathing glia promote axonal regeneration of rat retinal ganglion neurons. J Neurochem. 2003;85(4):861–71.

23. Pastrana E, Moreno-Flores MT, Avila J, Wandosell F, Minichiello L, Diaz-Nido J. BDNF production by olfactory ensheathing cells contributes to axonal regeneration of cultured adult CNS neurons. Neurochem Int. 2007;50(3):491–8.

24. Runyan SA, Phelps PE. Mouse olfactory ensheathing glia enhance axon outgrowth on a myelin substrate in vitro. Exp Neurol. 2009;216(1):95–104.

25. Woodhall E, West AK, Chuah MI. Cultured olfactory ensheathing cells express nerve growth factor, brain-derived neurotrophic factor, glia cell line-derived neurotrophic factor and their receptors. Brain Res Mol Brain Res. 2001;88(1-2):203–13.

26. Pastrana E, Moreno-Flores MT, Gurzov EN, Avila J, Wandosell F, Diaz-Nido J. Genes Associated with Adult Axon Regeneration Promoted by Olfactory Ensheathing Cells: A New Role for Matrix Metalloproteinase 2. J Neurosci. 2006;26(20):5347–59.

27. Simón D, Martín-Bermejo MJ, Gallego-Hernández MT, Pastrana E, García-Escudero V, García-Gómez A, et al. Expression of plasminogen activator inhibitor-1 by olfactory ensheathing glia promotes axonal regeneration. Glia. 2011;59(10):1458–71.

28. Asanuma H, Meldrum DR, Meldrum KK. Therapeutic applications of mesenchymal stem cells to repair kidney injury. J Urol. 2010;184(1):26–33.

29. Bianco P, Robey PG. Stem cells in tissue engineering. Nature. 2001;414(6859):118–21.

30. Chagastelles PC, Nardi NB, Camassola M. Biology and applications of mesenchymal stem cells. Sci Prog. 2010;93(Pt 2):113–27.

31. Pardal R, Clarke MF, Morrison SJ. Applying the principles of stem-cell biology to cancer. Nat Rev Cancer. 2003;3(12):895–902.

32. Czyz J, Wiese C, Rolletschek A, Blyszczuk P, Cross M, Wobus AM. Potential of embryonic and adult stem cells in vitro. Biol Chem. 2003;384(10-11):1391–409.

33. Delorme B, Chateauvieux S, Charbord P. The concept of mesenchymal stem cells. Regen Med. julio de 2006;1(4):497–509.

34. Ebert AD, Svendsen CN. Stem cell model of spinal muscular atrophy. Arch Neurol. 2010;67(6):665–9.

35. McCracken MN, George BM, Kao KS, Marjon KD, Raveh T, Weissman IL. Normal and Neoplastic Stem Cells. Cold Spring Harb Symp Quant Biol. 2016;81:1–9.

36. Zipori D. The stem state: plasticity is essential, whereas self-renewal and hierarchy are optional. Stem Cells. 2005;23(6):719–26.

37. Erceg S, Ronaghi M, Oria M, García Roselló M, Aragó MAP, Lopez MG, et al. Transplanted Oligodendrocytes and Motoneuron Progenitors Generated from Human Embryonic Stem Cells Promote Locomotor Recovery After Spinal Cord Transection. Stem Cells. 2010;28(9):1541–9.

38. Moreno-Manzano V, Rodríguez-Jiménez FJ, García-Roselló M, Laínez S, Erceg S, Calvo MT, et al. Activated spinal cord ependymal stem cells rescue neurological function. Stem Cells. 2009;27(3):733–43.

39. Thuret S, Moon LDF, Gage FH. Therapeutic interventions after spinal cord injury. Nat Rev Neurosci. 2006;7(8):628–43.

40. Namiot ED, Niemi JVL, Chubarev VN, Tarasov VV, Schiöth HB. Stem Cells in Clinical Trials on Neurological Disorders: Trends in Stem Cells Origins, Indications, and Status of the Clinical Trials. International Journal of Molecular Sciences. enero de 2022;23(19):11453.

41. Lu Y, Wang L, Zhang M, Chen Z. Mesenchymal Stem Cell-Derived Small Extracellular Vesicles: A Novel Approach for Kidney Disease Treatment. Int J Nanomedicine. 2022;17:3603–18.

42. Li M, Chen H, Zhu M. Mesenchymal stem cells for regenerative medicine in central nervous system. Frontiers in Neuroscience [Internet]. 2022 [citado 24 de octubre de 2023];16. Disponible en: https://www.frontiersin.org/articles/10.3389/fnins.2022.1068114

43. Li X, Ling W, Pennisi A, Wang Y, Khan S, Heidaran M, et al. Human placenta-derived adherent cells prevent bone loss, stimulate bone formation, and suppress growth of multiple myeloma in bone. Stem Cells. 2011;29(2):263–73.

44. Lloyd AC. Limits to lifespan. Nat Cell Biol. 2002;4(2):E25–27.

45. Chopp M, Zhang XH, Li Y, Wang L, Chen J, Lu D, et al. Spinal cord injury in rat: treatment with bone marrow stromal cell transplantation. Neuroreport. 2000;11(13):3001–5.

46. Hofstetter CP, Schwarz EJ, Hess D, Widenfalk J, El Manira A, Prockop DJ, et al. Marrow stromal cells form guiding strands in the injured spinal cord and promote recovery. Proc Natl Acad Sci U S A. 2002;99(4):2199–204.

47. Ankeny DP, McTigue DM, Jakeman LB. Bone marrow transplants provide tissue protection and directional guidance for axons after contusive spinal cord injury in rats. Exp Neurol. 2004;190(1):17–31.

48. Kamada T, Koda M, Dezawa M, Yoshinaga K, Hashimoto M, Koshizuka S, et al. Transplantation of Bone Marrow Stromal Cell-Derived Schwann Cells Promotes Axonal Regeneration and Functional Recovery after Complete Transection of Adult Rat Spinal Cord. J Neuropathol Exp Neurol. 2005;64(1):37–45.

49. Cizkova D, Novotna I, Slovinska L, Vanicky I, Jergova S, Rosocha J, et al. Repetitive intrathecal catheter delivery of bone marrow mesenchymal stromal cells improves functional recovery in a rat model of contusive spinal cord injury. J Neurotrauma. 2011;28(9):1951–61.

50. Himes BT, Neuhuber B, Coleman C, Kushner R, Swanger SA, Kopen GC, et al. Recovery of function following grafting of human bone marrow-derived stromal cells into the injured spinal cord. Neurorehabil Neural Repair. 2006;20(2):278–96.

51. Shichinohe H, Kuroda S, Tsuji S, Yamaguchi S, Yano S, Lee JB, et al. Bone Marrow Stromal Cells Promote Neurite Extension in Organotypic Spinal Cord Slice: Significance for Cell Transplantation Therapy. Neurorehabil Neural Repair. 2008;22(5):447–57.

52. Someya Y, Koda M, Dezawa M, Kadota T, Hashimoto M, Kamada T, et al. Reduction of cystic cavity, promotion of axonal regeneration and sparing, and functional recovery with transplanted bone marrow stromal cell-derived Schwann cells after contusion injury to the adult rat spinal cord. J Neurosurg Spine. 2008;9(6):600–10.

53. Zurita M, Vaquero J, Bonilla C, Santos M, De Haro J, Oya S, et al. Functional recovery of chronic paraplegic pigs after autologous transplantation of bone marrow stromal cells. Transplantation. 2008;86(6):845–53.

54. Li Y, Chen J, Chen XG, Wang L, Gautam SC, Xu YX, et al. Human marrow stromal cell therapy for stroke in rat: neurotrophins and functional recovery. Neurology. 2002;59(4):514–23.

55. Iihoshi S, Honmou O, Houkin K, Hashi K, Kocsis JD. A therapeutic window for intravenous administration of autologous bone marrow after cerebral ischemia in adult rats. Brain Res. 2004;1007(1-2):1–9.

56. Sasaki M, Radtke C, Tan AM, Zhao P, Hamada H, Houkin K, et al. BDNF-hypersecreting human mesenchymal stem cells promote functional recovery, axonal sprouting, and protection of corticospinal neurons after spinal cord injury. J Neurosci. 2009;29(47):14932–41.

57. Bonab MM, Alimoghaddam K, Talebian F, Ghaffari SH, Ghavamzadeh A, Nikbin B. Aging of mesenchymal stem cell in vitro. BMC Cell Biology. 2006;7(1):14.

58. Andrzejewska A, Dabrowska S, Lukomska B, Janowski M. Mesenchymal Stem Cells for Neurological Disorders. Advanced Science. 2021;8(7):2002944.

59. Moonshi SS, Adelnia H, Wu Y, Ta HT. Placenta-derived mesenchymal stem cells for treatment of diseases: a clinically relevant source. Advanced Therapeutics. 2022;5(10):1–14.

60. Samsonraj RM, Raghunath M, Nurcombe V, Hui JH, van Wijnen AJ, Cool SM. Concise Review: Multifaceted Characterization of Human Mesenchymal Stem Cells for Use in Regenerative Medicine. Stem Cells Transl Med. 2017;6(12):2173–85.

61. Trallori E, Ghelardini C, Di Cesare Mannelli L. Mesenchymal stem cells, implications for pain therapy. Neural Regen Res. 2019;14(11):1915–6.

62. Gu Q, Gu Y, Shi Q, Yang H. Hypoxia Promotes Osteogenesis of Human Placental-Derived Mesenchymal Stem Cells. Tohoku J Exp Med. 2016;239(4):287–96.

63. Haque N, Rahman MT, Abu Kasim NH, Alabsi AM. Hypoxic culture conditions as a solution for mesenchymal stem cell based regenerative therapy. ScientificWorldJournal. 2013;2013:632972.

64. Li L, Jaiswal PK, Makhoul G, Jurakhan R, Selvasandran K, Ridwan K, et al. Hypoxia modulates cell migration and proliferation in placenta-derived mesenchymal stem cells. J Thorac Cardiovasc Surg. 2017;154(2):543–552.e3.

65. Mohyeldin A, Garzón-Muvdi T, Quiñones-Hinojosa A. Oxygen in stem cell biology: a critical component of the stem cell niche. Cell Stem Cell. 2010;7(2):150–61.

66. Youssef A, Iosef C, Han VKM. Low-oxygen tension and IGF-I promote proliferation and multipotency of placental mesenchymal stem cells (PMSCs) from different gestations via distinct signaling pathways. Endocrinology. 2014;155(4):1386–97.

67. Zhang Y, Ma L, Su Y, Su L, Lan X, Wu D, et al. Hypoxia conditioning enhances neuroprotective effects of aged human bone marrow mesenchymal stem cell-derived conditioned medium against cerebral ischemia in vitro. Brain Res. 2019;1725:146432.

68. Zhu C, Yu J, Pan Q, Yang J, Hao G, Wang Y, et al. Hypoxia-inducible factor-2 alpha promotes the proliferation of human placenta-derived mesenchymal stem cells through the MAPK/ERK signaling pathway. Sci Rep. 2016;6(1):35489.

69. Plaza N, Simón D, Sierra J, Moreno-Flores MT. Transduction of an immortalized olfactory ensheathing glia cell line with the green fluorescent protein (GFP) gene: Evaluation of its neuroregenerative capacity as a proof of concept. Neuroscience Letters. 2016;612:25–31.

70. Sánchez Martin C, Ledesma D, Dotti CG, Avila J. Microtubule-associated protein-2 located in growth regions of rat hippocampal neurons is highly phosphorylated at its proline-rich region. Neuroscience. 2000;101(4):885–93.

71. M D, K LB, I M, I SC, F M, D K, et al. Minimal criteria for defining multipotent mesenchymal stromal cells. The International Society for Cellular Therapy position statement. Cytotherapy [Internet]. 2006 [citado 20 de octubre de 2023];8(4). Disponible en: https://pubmed.ncbi.nlm.nih.gov/16923606/

72. Tirpe AA, Gulei D, Ciortea SM, Crivii C, Berindan-Neagoe I. Hypoxia: Overview on Hypoxia-Mediated Mechanisms with a Focus on the Role of HIF Genes. International Journal of Molecular Sciences. 2019;20(24):6140.

73. Ciruelas K, Marcotulli D, Bajjalieh SM. Synaptic vesicle protein 2: A multi-faceted regulator of secretion. Seminars in Cell & Developmental Biology. 2019;95:130–41.

74. Nowack A, Yao J, Custer KL, Bajjalieh SM. SV2 regulates neurotransmitter release via multiple mechanisms. Am J Physiol Cell Physiol. 2010;299(5):C960–7.

75. Catterall WA, Kalume F, Oakley JC. NaV1.1 channels and epilepsy. J Physiol. 2010;588(Pt 11):1849–59.

76. Rhodes KJ, Carroll KI, Sung MA, Doliveira LC, Monaghan MM, Burke SL, et al. KChIPs and Kv4 alpha subunits as integral components of A-type potassium channels in mammalian brain. J Neurosci. 2004;24(36):7903–15.

77. Campbell A, Brieva T, Raviv L, Rowley J, Niss K, Brandwein H, et al. Concise Review: Process Development Considerations for Cell Therapy. Stem Cells Transl Med. 2015;4(10):1155–63.

78. Morizane A. Cell therapy for Parkinson’s disease with induced pluripotent stem cells. Rinsho Shinkeigaku. 2019;59(3):119–24.

79. Pisciotta A, Bertoni L, Vallarola A, Bertani G, Mecugni D, Carnevale G. Neural crest derived stem cells from dental pulp and tooth-associated stem cells for peripheral nerve regeneration. Neural Regen Res. 2020;15(3):373–81.

80. Paradisi M, Alviano F, Pirondi S, Lanzoni G, Fernandez M, Lizzo G, et al. Human mesenchymal stem cells produce bioactive neurotrophic factors: source, individual variability and differentiation issues. Int J Immunopathol Pharmacol. 2014;27(3):391–402.

81. Bravery CA, Carmen J, Fong T, Oprea W, Hoogendoorn KH, Woda J, et al. Potency assay development for cellular therapy products: an ISCT review of the requirements and experiences in the industry. Cytotherapy. 2013;15(1):9–19.

82. Samsonraj RM, Raghunath M, Nurcombe V, Hui JH, van Wijnen AJ, Cool SM. Concise Review: Multifaceted Characterization of Human Mesenchymal Stem Cells for Use in Regenerative Medicine. Stem Cells Transl Med. 2017;6(12):2173–85.

83. Zhou T, Yuan Z, Weng J, Pei D, Du X, He C, et al. Challenges and advances in clinical applications of mesenchymal stromal cells. Journal of Hematology & Oncology. 12 de febrero de 2021;14(1):24.

84. Shammaa R, El-Kadiry AEH, Abusarah J, Rafei M. Mesenchymal Stem Cells Beyond Regenerative Medicine. Frontiers in Cell and Developmental Biology [Internet]. 2020 [citado 25 de octubre de 2023];8. Disponible en: https://www.frontiersin.org/articles/10.3389/fcell.2020.00072

85. Hoang DM, Pham PT, Bach TQ, Ngo ATL, Nguyen QT, Phan TTK, et al. Stem cell-based therapy for human diseases. Sig Transduct Target Ther. 6 de agosto de 2022;7(1):1–41.

86. Vellasamy S, Sandrasaigaran P, Vidyadaran S, George E, Ramasamy R. Isolation and characterisation of mesenchymal stem cells derived from human placenta tissue. World Journal of Stem Cells. 2012;4(6):53.

87. Kong T, Park JM, Jang JH, Kim CY, Bae SH, Choi Y, et al. Immunomodulatory effect of CD200-positive human placenta-derived stem cells in the early phase of stroke. Exp Mol Med. 2018;50(1):e425–e425.

88. Belmar-Lopez C, Mendoza G, Oberg D, Burnet J, Simon C, Cervello I, et al. Tissue-derived mesenchymal stromal cells used as vehicles for anti-tumor therapy exert different in vivo effects on migration capacity and tumor growth. BMC Med. 2013;11:139.

89. Parolini O, Alviano F, Bagnara GP, Bilic G, Bühring HJ, Evangelista M, et al. Concise review: isolation and characterization of cells from human term placenta: outcome of the first international Workshop on Placenta Derived Stem Cells. Stem Cells. 2008;26(2):300–11.

90. Saleh M, Taher M, Sohrabpour AA, Vaezi AA, Nasiri Toosi M, Kavianpour M, et al. Perspective of placenta derived mesenchymal stem cells in acute liver failure. Cell Biosci. 2020;10:71.

91. Scheper V, Schwieger J, Hamm A, Lenarz T, Hoffmann A. BDNF-overexpressing human mesenchymal stem cells mediate increased neuronal protection in vitro. J Neurosci Res. 2019;97(11):1414–29.

92. Bari E, Perteghella S, Di Silvestre D, Sorlini M, Catenacci L, Sorrenti M, et al. Pilot Production of Mesenchymal Stem/Stromal Freeze-Dried Secretome for Cell-Free Regenerative Nanomedicine: A Validated GMP-Compliant Process. Cells. 2018;7(11):190.

93. Taghi GM, Ghasem Kashani Maryam H, Taghi L, Leili H, Leyla M. Characterization of in vitro cultured bone marrow and adipose tissue-derived mesenchymal stem cells and their ability to express neurotrophic factors. Cell Biol Int. 2012;36(12):1239–49.

94. Marconi S, Castiglione G, Turano E, Bissolotti G, Angiari S, Farinazzo A, et al. Human Adipose-Derived Mesenchymal Stem Cells Systemically Injected Promote Peripheral Nerve Regeneration in the Mouse Model of Sciatic Crush. Tissue Engineering Part A. 2012;18(11-12):1264–72.

95. Oses C, Olivares B, Ezquer M, Acosta C, Bosch P, Donoso M, et al. Preconditioning of adipose tissue-derived mesenchymal stem cells with deferoxamine increases the production of pro-angiogenic, neuroprotective and anti-inflammatory factors: Potential application in the treatment of diabetic neuropathy. PLoS One. 2017;12(5):e0178011.

96. Sonigra RJ, Brighton PC, Jacoby J, Hall S, Wigley CB. Adult rat olfactory nerve ensheathing cells are effective promoters of adult central nervous system neurite outgrowth in coculture. Glia. 1999;25(3):256–69.

97. Sakmann B, Neher E. Patch Clamp Techniques for Studying Ionic Channels in Excitable Membranes. Annual Review of Physiology. 1984;46(1):455–72.

